# TnCentral: A Prokaryotic Transposable Element Database and Web Portal for Transposon Analysis

**DOI:** 10.1101/2021.05.26.445724

**Authors:** Karen Ross, Alessandro M. Varani, Erik Snesrud, Hongzhan Huang, Danillo Oliveira Alvarenga, Jian Zhang, Cathy Wu, Patrick McGann, Mick Chandler

**Affiliations:** Protein Information Resource, Department of Biochemistry and Molecular & Cellular Biology, Georgetown University Medical Center, Washington, DC, USA; School of Agricultural and Veterinary Sciences, Universidade Estadual Paulista, Jaboticabal, Sao Paulo, Brazil; Multidrug-Resistant Organism Repository and Surveillance Network, Walter Reed Army Institute of Research, Silver Spring, Maryland, USA; Protein Information Resource, Center for Bioinformatics and Computational Biology, University of Delaware, USA; Department of Biochemistry and Molecular & Cellular Biology, Georgetown University Medical Center, Washington, DC, USA

**Keywords:** Mobile Genetic Elements, genome evolution, antibiotic resistance, virulence

## Abstract

We describe here the structure and organization of TnCentral (https://tncentral.proteininformationresource.org/). a web resource for prokaryotic transposable elements (TE). TnCentral currently contains ~400 carefully annotated TE, including transposons from the Tn*3*, Tn*7*, Tn*402* and Tn*554* families, compound transposons, integrons and associated insertion sequences (IS). These TE carry passenger genes, including genes conferring resistance to over 25 classes of antibiotics and nine types of heavy metal as well as genes responsible for pathogenesis in plants, toxin/antitoxin gene pairs, transcription factors and genes involved in metabolism. Each TE has its own entry page providing details about its transposition genes, passenger genes, and other sequence features required for transposition as well as a graphical map of all features. TnCentral content can be browsed and queried through text and sequence-based searches with a graphic output. We describe three use cases, which illustrate how the search interface, results tables, and entry pages can be used to explore and compare TEs.

TnCentral also includes downloadable software to facilitate user-driven identification, with manual annotation, of certain types of TE in genomic sequences. Through the TnCentral homepage, users can also access TnPedia which provides comprehensive reviews of the major TE families including an extensive general section, and specialised sections with descriptions of insertion sequence and transposon families. TnCentral and TnPedia are intuitive resources that can be used by clinicians and scientists to assess TE diversity in clinical, veterinary and environmental samples.

**Importance:** The ability of bacteria to undergo rapid evolution and adapt to changing environmental circumstances drives the public health crisis of multiple antibiotic resistance as well as outbreaks of disease in economically important agricultural crops and animal husbandry. Prokaryotic transposable elements (TE) play a critical role in this. Many carry “passenger genes” (not required for the transposition process) conferring resistance to antibiotics or heavy metals or causing disease in plants and animals. Passenger genes are spread by the normal TE transposition activities, by insertion into plasmids which then spread via conjugation within and across bacterial populations. Thus, an understanding of TE composition and transposition mechanisms is key to developing strategies to combat bacterial pathogenesis. Toward this end, we have developed TnCentral, a bioinformatics resource dedicated to describing and exploring the structural and functional features of prokaryotic TE and whose use is intuitive and accessible to users with or without bioinformatics expertise.

## Introduction

Transposable elements (TE) are key facilitators of bacterial evolution and adaptation and central players in the emergence of antibiotic and heavy metal resistance and to the transmission of virulence and pathogenic traits. Some TE can capture “passenger genes” (genes not involved in the transposition process) encoding these traits and transmit them to plasmids, where they accumulate and are then transferred within and between bacterial populations by conjugation. TE also contribute significantly to the on-going reorganization of bacterial genomes giving rise to new strains that are more adept at proliferating in clinical and agricultural environments, as well as in natural ecosystems.

Understanding TE nature, distribution, and activity is therefore an indispensable part of the struggle to cope with the public health crisis of multiple antibiotic resistance (ABR) [1,2]. To understand the impact of TE on bacterial populations, it is essential to provide a detailed description and catalog of TE structures and diversity. The simplest TE, known as Insertion Sequences (IS), have a profound impact on genome organization and function (see [3–7]) but do not themselves generally carry integrated passenger genes. There are a large number of significantly more complex TE (Figure 1), arguably even more important in the global emergence of ABR and other virulence and pathogenicity traits. These are generically called transposons and may carry multiple passenger genes, including some of the most clinically important antibiotic resistance genes. Like IS, these TE are grouped into a number of distinct families with characteristic organizations [3]. Their transposition activities facilitate the rapid spread of groups of antibiotic resistance genes and promote their horizontal transfer. Yet another important aspect of their impact is their ability to assemble passenger genes into resistance clusters [8,9]. While there appears to be a wide-spread appreciation that mobile plasmids are responsible for the spread of antibiotic resistance, it is less well-known that IS and transposons are the conduits that transfer this information between chromosomes and plasmids.

**Figure 1.**
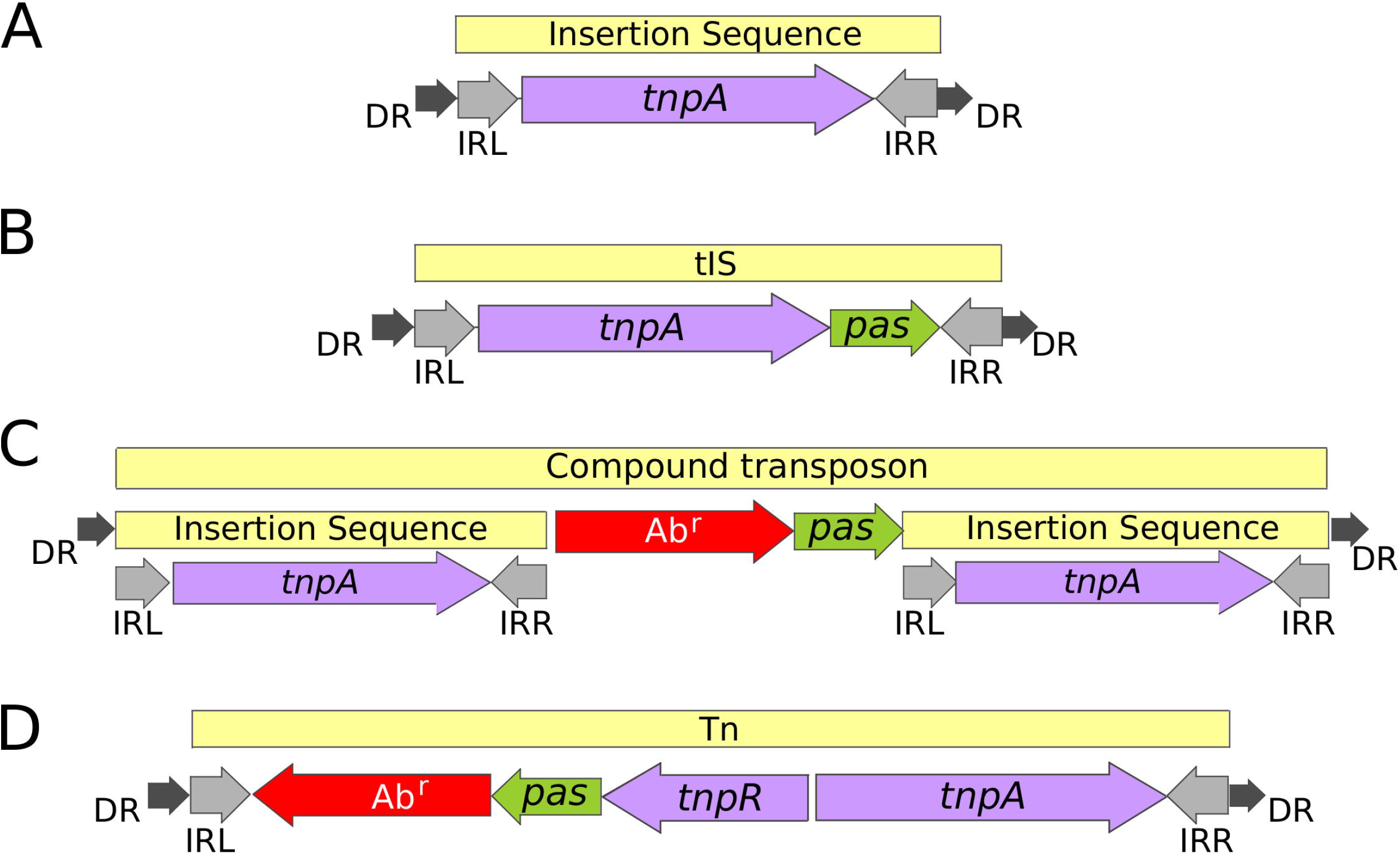
Structural arrangement of prokaryotic Transposable Elements. The TE is indicated by a pale-yellow horizontal bar at the top of each section. Open reading frames are shown as horizontal arrows with the arrowheads indicating the direction of expression: purple, transposition-associated genes; red, antibiotic resistance genes; green, other passenger genes. The inverted terminal repeats found at the ends of the majority of TE are shown as grey arrows and the direct target repeats generally produced by insertion are indicated by small black arrows. **A.** Insertion Sequence (IS), a short DNA segment encoding only the mobilization protein (Transposase, TnpA), flanked by two imperfect Inverted Repeats (IRs), and generally containing a short flanking directly repeated duplication (DR) on the target of insertion. **B.** tIS (transporter IS) are structurally similar to an IS, but contain passenger genes. They are presently restricted to the IS*1595* and IS*66* families. **C.** Compound transposons are formed by two IS in either direct or inverted orientation, flanking a variety of passenger genes including those for antibiotic resistance. **D.** Transposons are more heterogeneous structures and include different sets of transposition-related genes which are specific to each Tn family and multiple antibiotic resistances, virulence and other passenger genes. This is an example of a Tn*3* family transposon with transposon, *tnpA*, and resolvase genes, *tnpR*.

There are a number of other bioinformatics resources that cover aspects of prokaryotic TE biology. These include databases for TE passenger genes such as antibiotic resistance (CARD [10], ARDB [11]) or toxin/antitoxin gene pairs (TADB [12], TASmania [13]) as well as the various classes of TE themselves such as insertion sequences (ISfinder [14]), integrons (INTEGRALL [15]), integrative conjugative elements, ICE (ICEberg [16,17]), plasmids (PlasmidFinder [18]) or more general databases which include a variety of these genome components (ACLAME [19–21]). However, there is a need for a resource that collects, compares and collates detailed information on the various different classes of TE that are responsible for the transmission of medically and economically important passenger genes in an intuitive and accessible way.

Here, we describe TnCentral (https://tncentral.proteininformationresource.org/), a database of detailed structural and functional information on bacterial TE. Additionally, TnCentral provides access to TnPedia (https://tnpedia.fcav.unesp.br/), a comprehensive encyclopedia describing the current state of our knowledge of the biology of IS and transposons. Together, TnCentral and TnPedia provide a detailed description of TE diversity with easy-to-understand graphics outputs that are accessible to users without significant bioinformatic knowledge. They allow users to rapidly analyse the landscape of TE in genomes (chromosomes and plasmids) isolated from clinical, veterinary and environmental samples.

## Results

### TnCentral Website Content

As of May 2021, TnCentral contains information on ~400 TE. About half of these TE are *Tn3*-family transposons. The remainder are integrons, compound transposons, transposons from the *Tn402, Tn554* and *Tn7* families, and IS that are associated with TE or are part of compound transposons (Supplementary Table 1). They include TE with resistance to over 25 different classes of antibiotics and nine different heavy metals. The collection also contains TE that carry a toxin/antitoxin system for bacterial plasmid maintenance [22–24] and TE from xanthomonads carrying genes for plant pathogenicity.

### TnCental Web Portal

The TnCentral home page is designed to give the user easy access to the contents of TnCentral with a number of options (Figure 2A), including:

**TnCentral Search** (*search of the TnCentral database*),
**Sequence Search** (*BLAST-like search for sequence similarities in the database*),
**Browse Tn list** (*view all TE in TnCentral*),
**Tnfinder Software** (*access to downloadable scripts for identifying potential TE in sequence databases*),
**Documentation** (*downloadable documentation for TnCentral*),
**For Curators** (*detailed curation guidelines*),
**TnPedia** (*TE Encyclopedia*),
**Related links**, *and*
**Feedback.**

**Figure 2.**
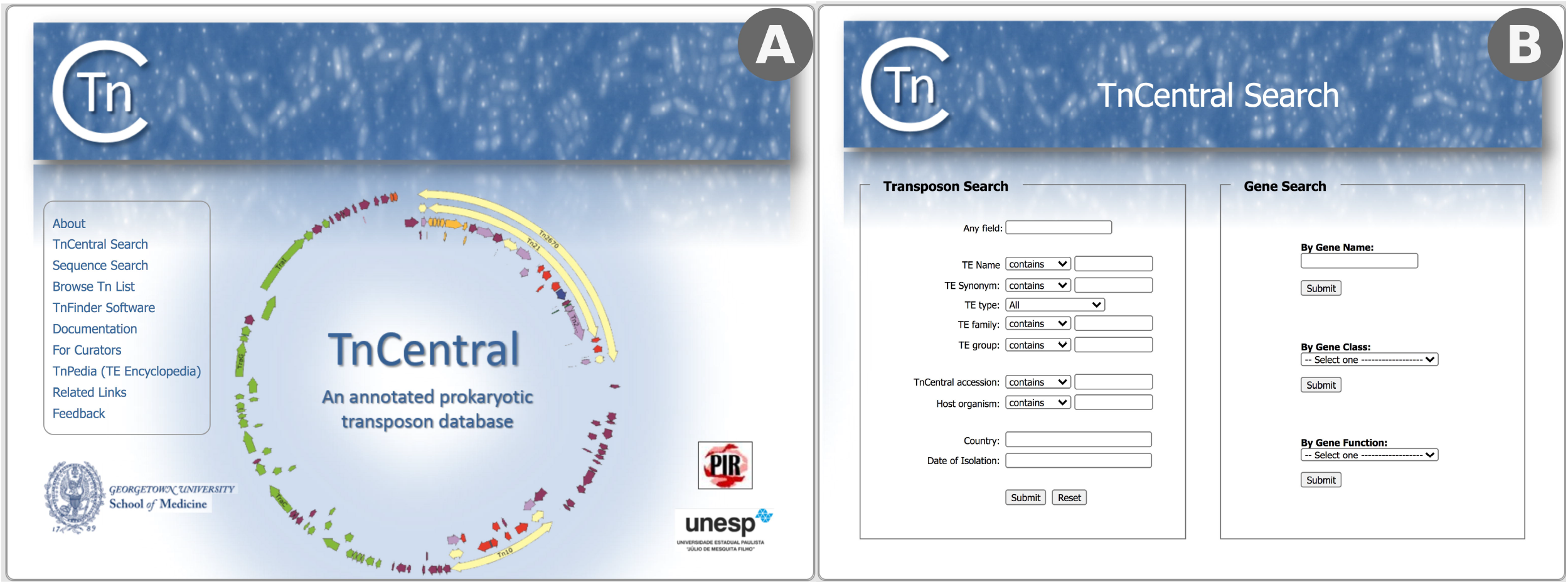
A) TnCentral homepage showing clickable links to various TnCentral sections in the box on the left. B) TnCentral search interface showing search choices for TE on the left and for transposition-related and passenger genes on the left.

#### TnCentral Search

The interface provides a variety of search functions divided into two search types: ***Transposon search*** and ***Gene search*** (Figure 2B).

##### Transposon search

The transposon collection can be searched using the transposon **name, synonyms** which may have been used in the literature, the **type** of mobile genetic element (e.g., insertion sequence, transposon or integron), the **family** and **subgroup** to which it belongs, the **host organism, country** of identification and **date** of identification. The latter three search terms are intended for use in epidemiological tracking. These search terms result in a table that can be sorted, customized and downloaded (See Use Case #1, below).

##### Gene search

It is also possible to search for TE-associated genes by name, by class (Transposase, Accessory Gene or Passenger Gene) or by function (Antibiotic Resistance, Heavy Metal Resistance) and to retrieve information on the transposons in which they are found (see Use Case #2, below).

##### Sequence Search

Sequence Search allows users to perform sequence similarity searches using BLAST [26,27] (see Use Case #3, below). By default, the search database is the TnCentral database, but the page also provides links to BLAST against the ISfinder (https://isfinder.biotoul.fr/blast.php), NCBI https://blast.ncbi.nlm.nih.gov/Blast.cgi), Comprehensive Antibiotic Resistance Database (CARD; https://card.mcmaster.ca/analyze/blast) and the Toxin-Anitoxin (TADB; https://bioinfo-mml.sjtu.edu.cn/TADB2/) databases. The BLAST tool automatically distinguishes between DNA and protein query sequences.

#### Browse Tn list

This option allows the user to browse the entire TnCentral database.

#### The Transposon entry page

All of the search and browse options provide links to entry pages for each TE (Figure 3), which provide detailed information about TE features and origins. The page includes: 1) host information: host species, strain, and plasmid/chromosome in which the transposon was found as well as the date and geographic location of the isolate; 2) a graphic representation of the annotated sequence with color-coded features; 3) Terminal Inverted Repeats (IR); 4) DNA sequence; 5) internal recombination sites (e.g. *res* sites) including their coordinates, length and DNA sequence; 6) ORF summary, which includes all protein coding genes in the order in which they appear, 5’-3’, in the TE sequence, the element with which they are associated (important for nested TE in which one TE is inserted into another), their coordinates, their class (e.g., Transposase, Accessory Gene, Passenger Gene) and subclass (e.g., Antibiotic Resistance, Heavy Metal Resistance) and their relative orientation within the TE; 7) a detailed ORF description including the amino acid sequence; 8) if applicable, a table of Internal Transposable Elements (TE inserted in the main element) including the name, type location and length; 9) if applicable, a table of Internal Repeats (repeat elements, other than the terminal inverted repeats, that are found within the TE), including the associated TE, coordinates and DNA sequence; 10) Bibliographic references with direct links to PubMed [25]. Each section can be collapsed using a button on the right-hand side of the section heading. Sections can be viewed either by scrolling down on the page or by clicking on the section name in the menu located on the left side of the page. Sequence files in FASTA and GenBank format can be downloaded using the links on the left side of the page under the menu.

**Figure 3.**
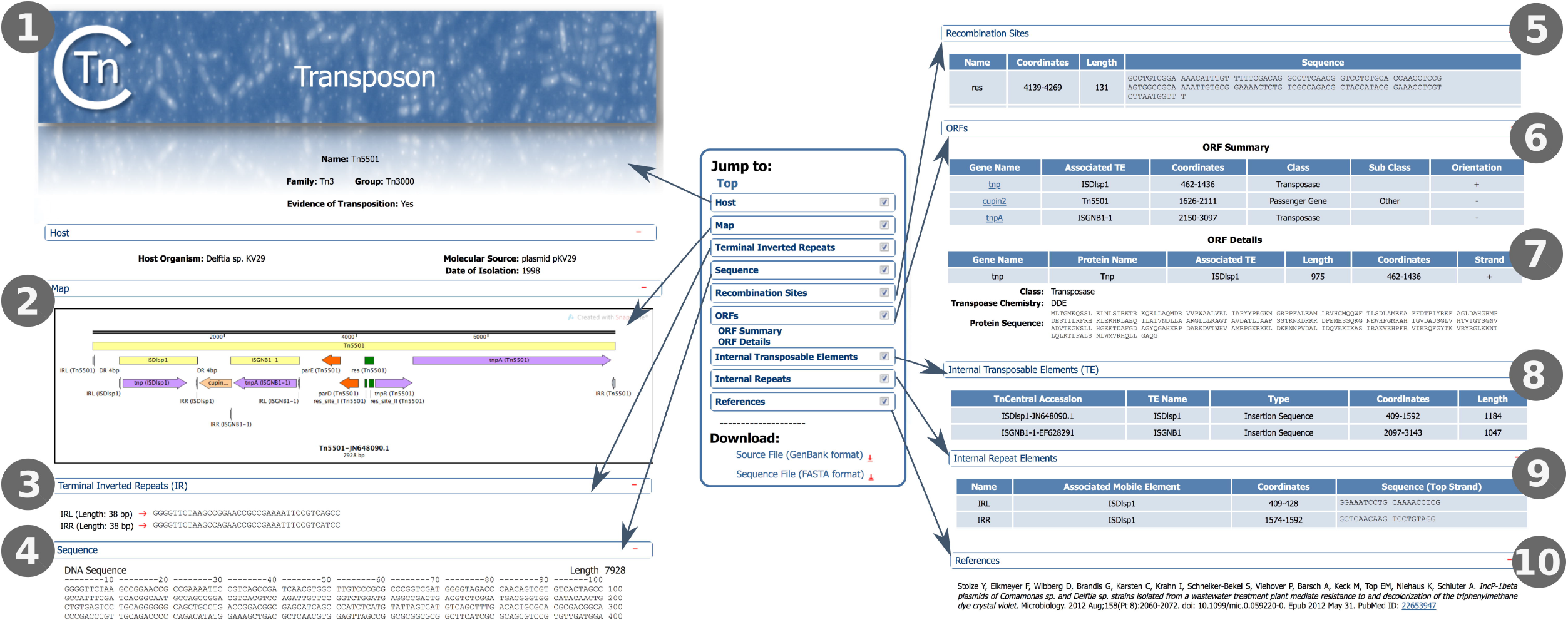
TnCentral TE Entry Page. #1-10: Sections of the entry page (see text for details).

#### Tnfinder Software

This section provides three user-downloadable scripts written in-house for identifying transposons. They provide users with local control over analyses and help them screen datasets containing large numbers of genomic sequences using their own servers for identifying potential candidates and which are then manually curated.

##### Tn3 Transposon Finder

(Tn3_finder) performs the automatic prediction of transposable elements of the Tn*3* family in bacteria and archaea. It compares user-provided bacterial and archaeal genome sequences to custom Tn*3* transposase and resolvase databases by BLAST alignments. The criteria for identifying potential transposon regions according to similarity, coverage and distance values can be adjusted by the user. Additional ORFs that might be related to passenger genes are also predicted, and flanking regions can also be retrieved and analyzed. The automatic prediction results are written in report files and pre-annotated GenBank files to help in subsequent manual curation. Tn*3*_finder allows for the concurrent analysis of multiple genomes by multithreading.

##### Composite Transposon Finder

(TnComp_finder) predicts the putative composite transposons in bacterial and archaeal genomes based on insertion sequence replicas in a relatively short span. It works by comparing nucleotide sequences from bacterial and archaeal genomes to a custom transposon database and identifying duplicated transposons in user-defined genomic regions from BLAST alignments. Similar to Tn3_finder, multithreaded analyses of multiple genomes are available and parameters for similarity, coverage, distance and flanking regions can be adjusted by the user. Results are written in report files and pre-annotated GenBank files to help in subsequent manual curation.

##### Antibiotic Resistance Gene-associated IS Finder

(ISAbR_finder) is an experimental program for the automatic prediction of antibiotic resistance genes associated with known IS elements derived from the ISfinder database and has yet to be tested extensively. It works by comparing IS nucleotide sequences from bacterial and archaeal genomes to a custom antibiotic resistance database based on the parsing of BLAST alignment results, using a number of parameters that can be customized by the user for stricter or more relaxed criteria and allowing multithreaded alignments of multiple genomes. ISAbR_finder also produces report files and pre-annotated GenBank files on which the recommended manual curation should be performed.

#### Documentation

This section, which can be downloaded as a pdf file, provides a short background description of transposons and TnCentral together with a short description of the **curation workflow** and of planned future developments.

#### For Curators

This section provides a detailed description of the curation workflow used in generating the annotated TnCentral data.

#### TnPedia

TnCentral provides access from the homepage to TnPedia, an online knowledge base which contains information concerning transposition in prokaryotes. TnPedia is developed using MediaWiki (https://www.mediawiki.org) and can also be accessed directly (https://tnpedia.fcav.unesp.br/). It is structured into three main sections: **General Information, IS Families and Transposon Families** (Figure 4).

**Figure 4.**
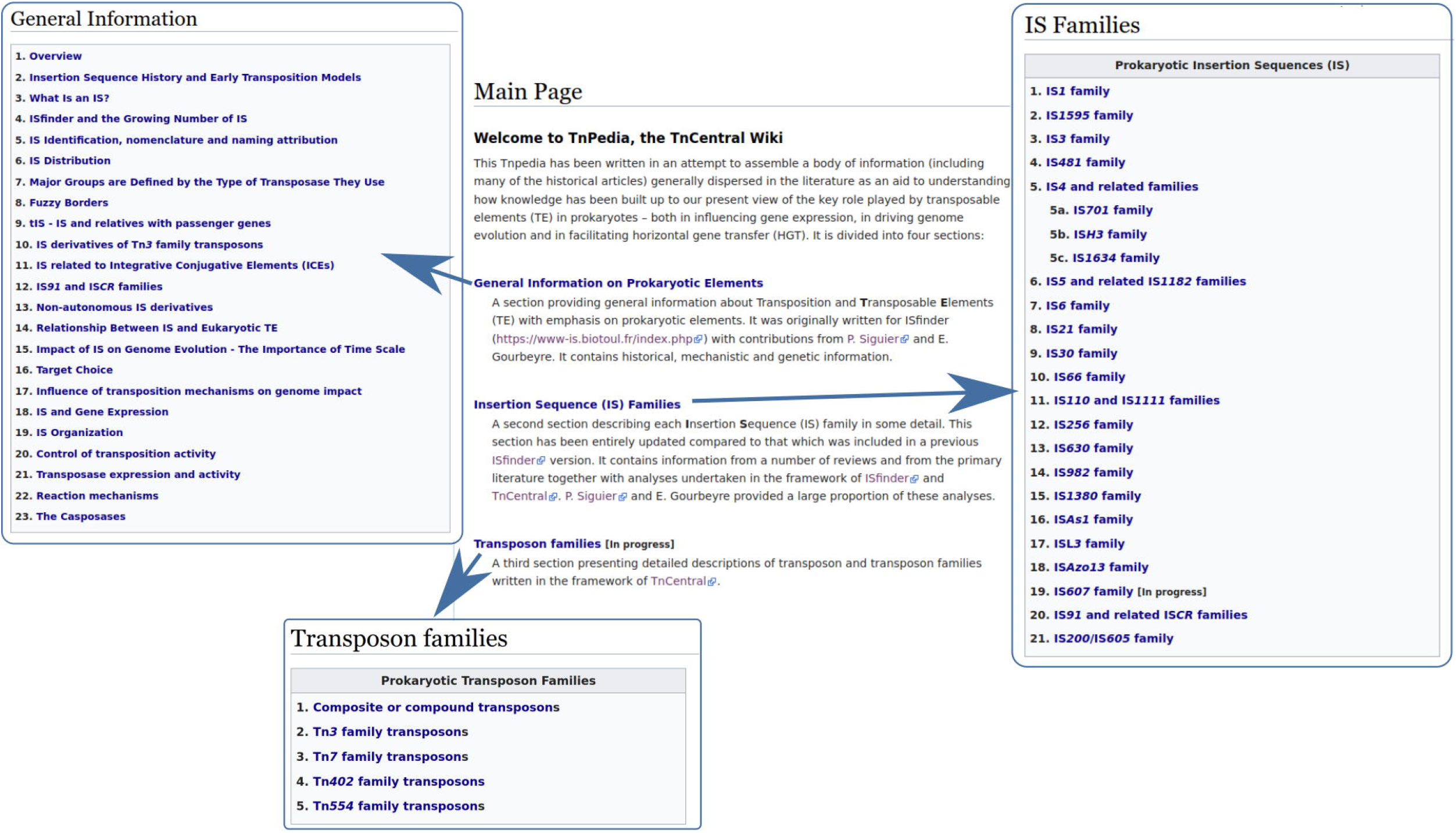
The main sections of TnPedia, a TnCentral-related wiki compiling information on prokaryotic transposable elements. Only three of the four sections (General Information, IS families and Transposon families) are illustrated. The fourth section is a Transposition Glossary, which is under construction.

The **General Information** section provides a series of clickable sections with an extensive bibliography and direct links to the articles in PubMed. It includes a historical perspective, definitions and descriptions of a variety of prokaryotic TE, the basic mechanisms involved in their movement and the enzymes involved in these processes. It also contains information describing their impact on their host genomes and how their activities are controlled.

The **IS Families** section consists of individual chapters describing each of the ~25 IS families in detail and covers, where possible, the identification of the founding members, their organisation, distribution, variability and phylogenetic relationships, regulation of their transposition, impact on their host genomes, and their transposition mechanisms including genetic, biochemical and structural studies.

The **Transposon Families** section describes each transposon family with similar information to that included in the IS family descriptions but, in addition, including a detailed description of their structures and the passenger genes which they may carry.

### Examples of TnCentral Use

#### Use Case #1: Comparing Protein Coding Genes in Tn*554* Family Members

The Tn*554* family is a small family restricted to the Firmicutes. Members encode three genes, *tnpA, tnpB* and *tnpC*, involved in transposition [28,29] (https://tnpedia.fcav.unesp.br/index.php/Transposons_families/Tn554_family). TnpA and TnpB both exhibit a C-terminal motif which shares all the important catalytic residues of a typical tyrosine site-specific recombinase [28,29]. They insert in a sequence-specific way into the DNA repair gene *radC* [30,31] and can also be found in a circular form [32–36]. To compare the protein coding genes in Tn*554* family members side by side, we searched for Tn*554* in the TE family field of the Transposon Search interface (Figure 5A). Fourteen Tn*554* family members were found (of which only 10 are shown in Figure 5B). In order to perform a side-by-side comparison of the protein-coding genes in these TE, we used the Customize Display option on the search results page, to add the “All Gene Fields” columns, which provide information about the protein coding genes, to the display and to remove several columns (e.g., Host Organism, Country) (Figure 5B). Results for two of the Tn*554* transposons (Tn*558.3* and Tn*559*) are shown in Figure 5C. Both transposons have the three-part transposition module (*tnpA, tnpB, tnpC*) characteristic of the family. However, the two transposons are quite diverse in their passenger genes. Tn*558.3* has gene called *fla*, which contains a flavodoxin-like domain, and the ABR gene *fexA*, which confers resistance to phenicol antibiotics. Tn*559* has just a single passenger gene, the ABR gene, *dfrK*, which confers resistance to diaminopyrimidine antibiotics. As shown by this example, the flexible search results page makes it easy to compare features across multiple transposons.

**Figure 5.**
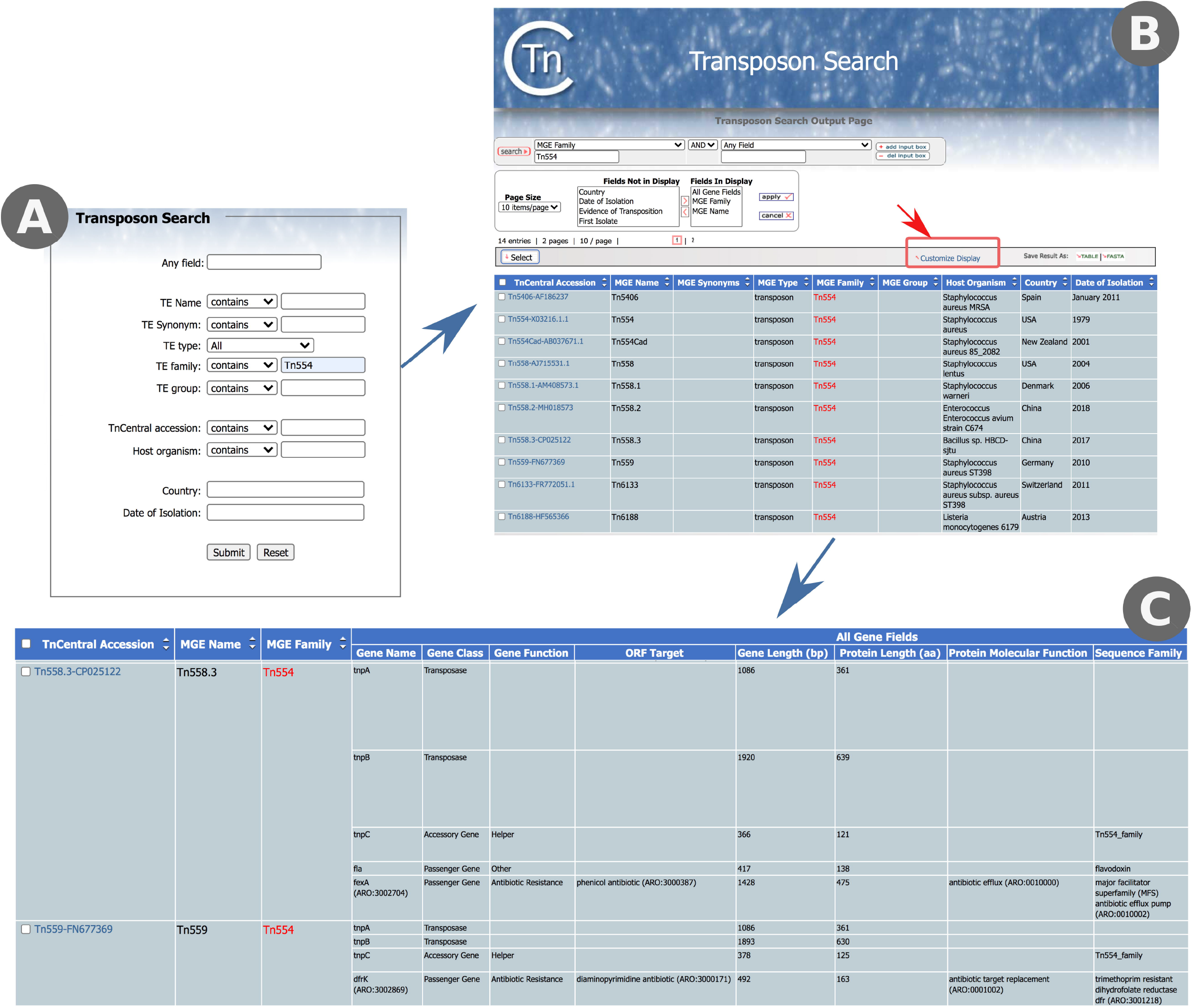
Comparing Protein Coding Genes in Tn*554* Family Members. A) TnCentral Transposon Search interface, showing a search for Tn*554* in the TE family field. B) Interface for customizing the columns in the search results display. Clicking on “Customize Display” (red box) opens the interface. C) Partial Tn*554* family search results after customization to show information on protein coding genes (All Gene Fields).

#### Use Case #2: Type II Toxin/Antitoxin Systems in Tn*3* Transposons

Toxin/Antitoxin (TA) systems are implicated in plasmid maintenance in bacterial populations [37]. These systems are characterized by a stable toxin and an unstable antitoxin that binds to the toxin and inhibits its lethal effect. Loss of a plasmid carrying a TA system will lead to rapid depletion of the antitoxin, allowing the persistent toxin to kill the cell. Thus, only members of a population that retain the plasmid will survive. Recently, a set of Tn*3*-family transposons carrying TA systems were characterized and included in the TnCentral database [22]. To explore these Tn, we used the TnCentral Gene Search function, selecting “Passenger Gene” from the Gene Class pull-down menu and “Toxin” from the Gene Sub-Class pull-down menu (Figure 6A, red box). The search results included eight different toxin genes (Gp49, HEPN, PIN, PIN_3, *abiEii, higB, parE*, and zeta) found in 43 different transposons. Similarly, transposons carrying antitoxin genes were identified using the Gene Search function with the Gene Sub-Class menu set to “Antitoxin” (Figure 6B, red box). There were 44 transposons carrying 11 different antitoxin genes. Combinations of toxin and antitoxin genes in individual transposons were examined by going to the ORF Summary section of the entry pages for the TA transposons. For example, Tn*Sku1* (Figure 6B, yellow box; Figure 6C) has a Gp49 toxin gene and an antitoxin gene containing an HTH domain (referred to as HTH). Most transposons have a single toxin/antitoxin gene pair except for Tn*Xca1*, which has two TA pairs, and Tn*5501.5*, which has a *parD* antitoxin gene and no toxin gene. The majority of Tn*5501* derivatives in TnCentral have a *parE* toxin gene as well as the *parD* antitoxin, suggesting that Tn*5501.5* may have undergone a deletion in the region containing *parE* (Supplementary Figure 1).

**Figure 6.**
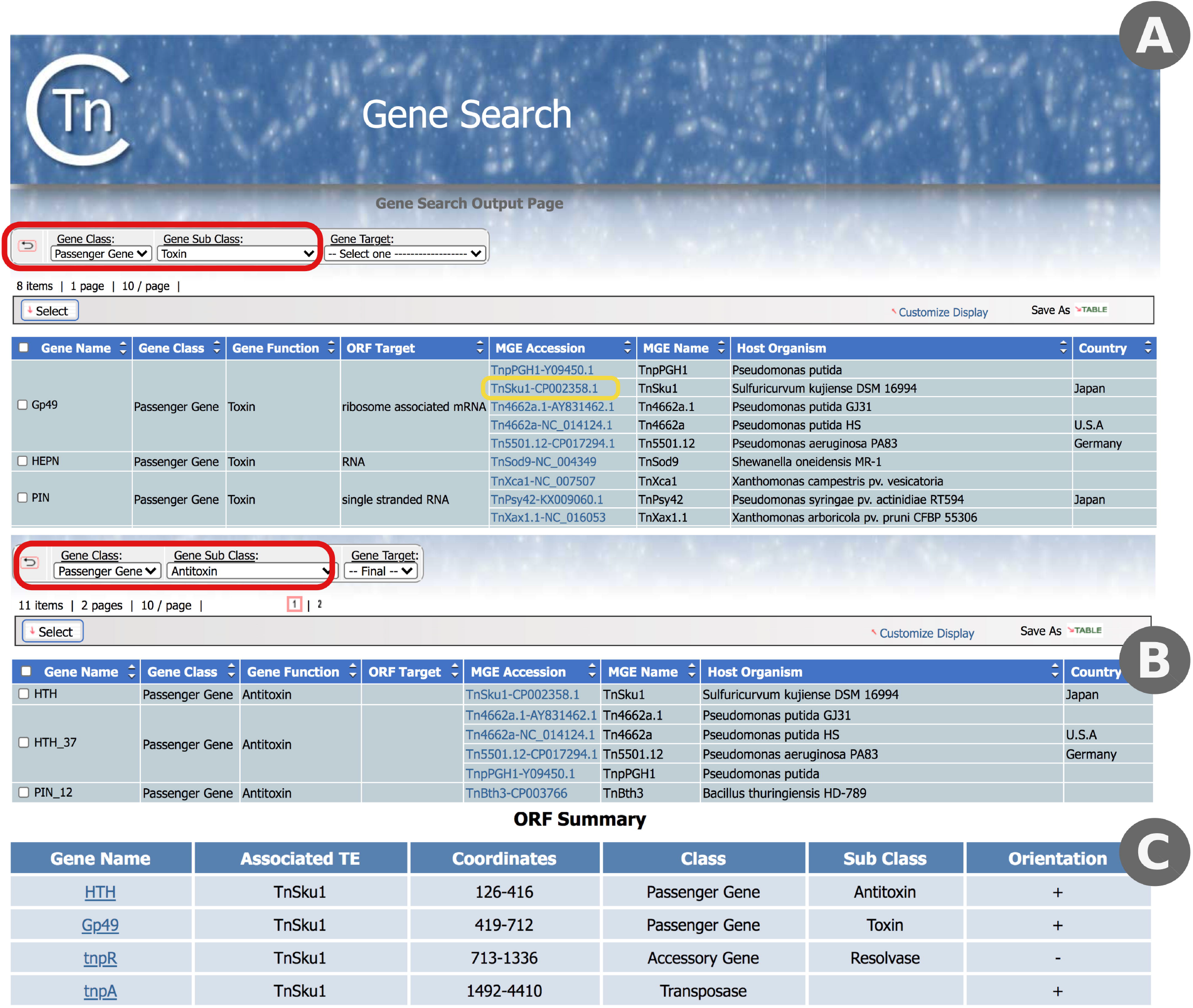
Exploring Toxin/Antitoxin Genes in TnCentral. A) Partial results of searching TnCentral for toxin genes. The settings used to obtain these results are shown in the red box. Links to entry pages for the TE carrying the indicated genes are provided in the MGE Accession column (e.g., Tn*Sku1*-CP002358.1, yellow box). B) Partial search results for antitoxin genes in TnCentral. Settings are shown in the red box. C) ORF Summary section of the entry page for Tn*Sku1*-CP002358.1, showing the presence of a toxin/antitoxin gene pair (Gp49 toxin/HTH antitoxin).

#### Use Case #3: Tn*21* and its Relatives

Tn*21* is the canonical member of a subfamily of Tn*3* transposons that confers a variety of antibiotic resistances [38–40] and several analyses have proposed mechanisms to explain how *Tn21* arose from simpler ancestor transposons (e.g., [40,41]). Tn*21* has a mercury resistance operon at the 5’- (left) end, a *tnpA/tnpR* transposition module at the 3’-(right) end, and a transposition-deficient integron (In2) carrying several ABR genes (a GCN5-related N-acetyltransferase (GNAT_fam), *sul1, qacEdelta1*, and *aadA*) in the middle (Supplementary Figure 2). These ABR genes confer resistance to aminoglycosides, sulfones, sulfonamides, quaternary ammonium salts, and acridine dye. More recently, a transposon that lacks the integron insertion but is otherwise identical to Tn*21* (the hypothetical Tn*21* backbone Tn21Δ in [40]) was discovered [42]. This transposon, Tn*5060*, was proposed to be the ancestor of Tn*21* [42]. Tn*21* also has numerous relatives that carry different combinations of antibiotic resistance genes within and outside the integron. To explore the Tn*21* subfamily, we performed a TnCentral Sequence Search (BLAST) using the putative ancestral Tn*5060* sequence (Figure 7A). In addition to Tn*5060* itself, we identified ten transposons in the database (Tn*20*, Tn*21*, Tn*21.1*, Tn*21.2*, Tn*5086*, Tn*2411*, Tn*2424*, Tn*4*, Tn*1935*, and Tn*As3*; Supplementary Figure 2) that contain all (or nearly all) of the Tn*5060* sequence. With the exception of Tn*20*, which is almost identical to Tn*5060* (99.5%), these transposons have two or more discontinuous sub-regions that align to Tn*5060*. For example, Tn*21* has two sub-regions, one of which is a close match to the left half of Tn*5060* and the other of which is a close match to the right half of Tn*5060* (red bars in Figure 7B). This suggests that these transposons arose from Tn*5060* via the insertion of other sequences.

**Figure 7.**
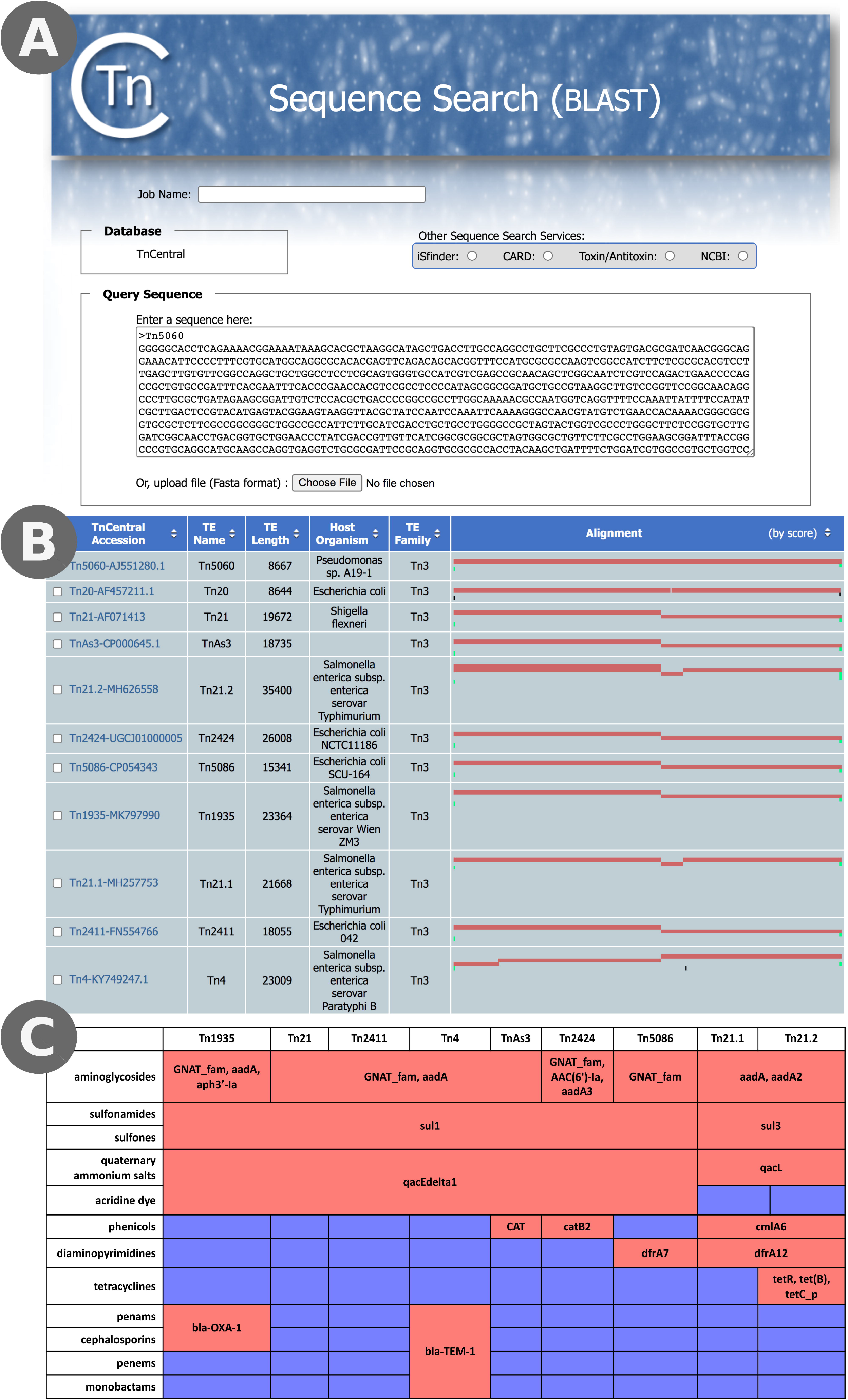
Analysis of ABR in Tn*21* Relatives. A) TnCentral Sequence Search using the sequence of Tn*5060*, the proposed ancestor of Tn*21*, as a query. B) Sequence Search results. The query sequence is represented by the width of the Alignment column. The red bars represent regions of the matched transposons that are highly similar to regions of Tn*5060*. C) ABR genes and targeted antibiotic classes in Tn*21* relatives. Red shading in the table cells indicates that the transposon carries at least one gene targeting the antibiotic class; blue shading indicates that it does not. The ABR genes found in each transposon are indicated in the table cells.

We compared the antibiotic resistance profiles of the ten transposons by inspecting their TnCentral entry pages. Tn*20*, like Tn*5060*, carries no ABR genes. The other nine transposons carry ABR genes targeting aminoglycosides, sulfones, sulfonamides, and quaternary ammonium salts (Figure 7C). Other resistances found in a subset of the six include acridine dye (Tn*1935*, Tn*21*, Tn*2411*, Tn*4*, Tn*As3*, Tn*2424*, Tn*5086*), carbapenams (Tn*1935* and Tn*4*), cephalosporins (Tn*1935* and Tn*4*), carbapenems (Tn*4*), monobactams (Tn*4*), phenicols (Tn*As3*, Tn*2424*, Tn*21.1*, Tn*21.2*), diaminopyrimidines (Tn*5086*, Tn*21.1*, Tn*21.2*), and tetracyclines (Tn*21.2*). Interestingly, in some cases where the transposons have resistances in common, they are conferred by different genes (Figure 7C). For example, phenicol resistance is conferred by *CAT* in Tn*As3, catB2 in* Tn*2424*, and *cmlA6* in Tn*21.1* and Tn*21.2*. Similarly, sulfonamide and sulfone resistance is conferred by *sul1* in all of the antibiotic-resistant family members except for Tn*21.1* and Tn*21.2*, where those resistances are conferred by *sul3*. Thus, even this closely related subfamily of transposons shows diversity in its antibiotic resistance genes. This is partially due to the flexibility of the integron to incorporate new antibiotic resistance gene cassettes but also to insertion of ABR-gene containing elements outside of the integron region (e.g., Tn*3.1* in Tn*4*, Supplementary Figure 2).

## Discussion

Here, we have described TnCentral, a user-friendly resource for exploration of prokaryotic TE. TnCentral provides a flexible search interface, TE-specific entry pages with intuitive graphics and detailed information about TE features, and a BLAST interface that allows users to identify TE that carry features of interest or to identify TE that are present in sequences of interest (e.g., plasmids). As shown in the use cases, the flexible search results page makes it easy to compare features across multiple transposons, the detailed entry pages allow exploration of TE passenger genes, such as ABR genes, and the Sequence Search enables retrieval of TE with related sequences that could be used as a starting point for evolutionary analyses. Moreover, TnCentral provides access to Tnfinder software for locating candidate TE in sequence data and to TnPedia, a comprehensive review of the biology of selected TE families.

As discussed in the Introduction, a variety of resources dedicated to aspects of prokaryotic TE biology currently exist. TnCentral’s unique contribution to this universe of resources lies in its coverage of a variety of TE (e.g. different transposon families and compound transposons with their associated IS and integrons) and its detailed focus on both core transposition genes and passenger genes of clinical, environmental, and economic importance. It has the additional feature of providing a clear graphic output for visualizing the often complex structures of TE.

The next step beyond annotation of individual TE is to annotate and visualize the TE content of prokaryotic chromosomes and plasmids. These studies are critical for understanding the propagation of high impact passenger genes, such as those that confer antibiotic resistance. Several tools that address this problem are available. For example, ISsaga [43], which is integrated into ISfinder, annotates IS present in user-provided sequences. Other software suites have been designed specifically to annotate IS in short read raw data (e.g. ISQuest [44], Transposon Insertion Finder [45], ISMapper [46] and panISa [47]) using preassembled libraries of TE and their components, while yet other approaches are based on *ab initio* prediction (e.g., OASIS [48], ISseeker [49] ISEscan [50], or provide a comparative view of IS mobilisation events (e.g. ISCompare [51]). These annotation tools are only as good as their underlying TE databases. ISfinder, which includes nearly 6000 individual examples of IS classified in distinct families and subfamilies according to their transposition mechanism and structural organization, provides such a rigorous framework for IS and has been incorporated into a number of annotation pipelines (e.g., ISsaga [43], MobileElementFinder [52]). However, IS represent only a fraction of prokaryotic TE, and unlike transposons and integrons, they rarely carry passenger genes. We hope that TnCentral will become a benchmark for more complex TE as ISfinder is for IS.

TnCentral is an ongoing project, and we will continue to expand and update the content. In addition to the exporting annotated TE in GenBank format, we plan to make all files available in a SnapGene file format which will allow users to use SnapGene (https://www.snapgene.com/), a commercial software tool (with a free viewer version) for visualizing and documenting nucleotide sequences and their features, to analyze and explore them. We also intend to enhance the visualization of TnCentral Sequence Search (i.e., BLAST) results to better support the analysis of plasmid sequences that may carry a complex complement of TE although it should be noted that the Sequence Search tool can already accommodate analysis of large plasmids. Ultimately, we envision that TnCentral could be used to analyze the TE content of a collection of sequences, such as patient, veterinary and environmental samples from an antibiotic resistance outbreak, to understand TE-driven evolution of the prokaryotic mobilome.

## Methods

### Curation Workflow

The TnCentral curation workflow is depicted in Figure 8. Curation is performed by members of the TnCentral development team as well as by graduate students in bioinformatics courses at Georgetown University Medical Center. TnFinder scripts are run against RefSeq and other sequence databases and GenBank files potentially containing TE are retrieved. TE sequences are isolated and annotated using SnapGene (https://www.snapgene.com). Features of interest (i.e., protein coding genes, TE, repeat elements, and recombination sites) are annotated according to detailed curation guidelines (provided in the “For Curators” of TnCentral). Fully annotated features are saved in a SnapGene Custom Library. New transposon sequences can be searched against this library, enabling detection of features previously identified in other TEs. All annotated TE files are checked by a second curator. An enhanced GenBank file containing all annotations is exported from SnapGene and checked for common curation formatting errors using a custom Perl script. Detected errors are manually corrected in the SnapGene file, which is then exported as a revised enhanced GenBank file. Information from this GenBank file is used to populate the TnCentral database, which, in turn, serves as the backend for the TnCentral web portal. An image file showing a color-coded map of TE features is also exported from SnapGene and displayed on the TE entry page.

**Figure 8.**
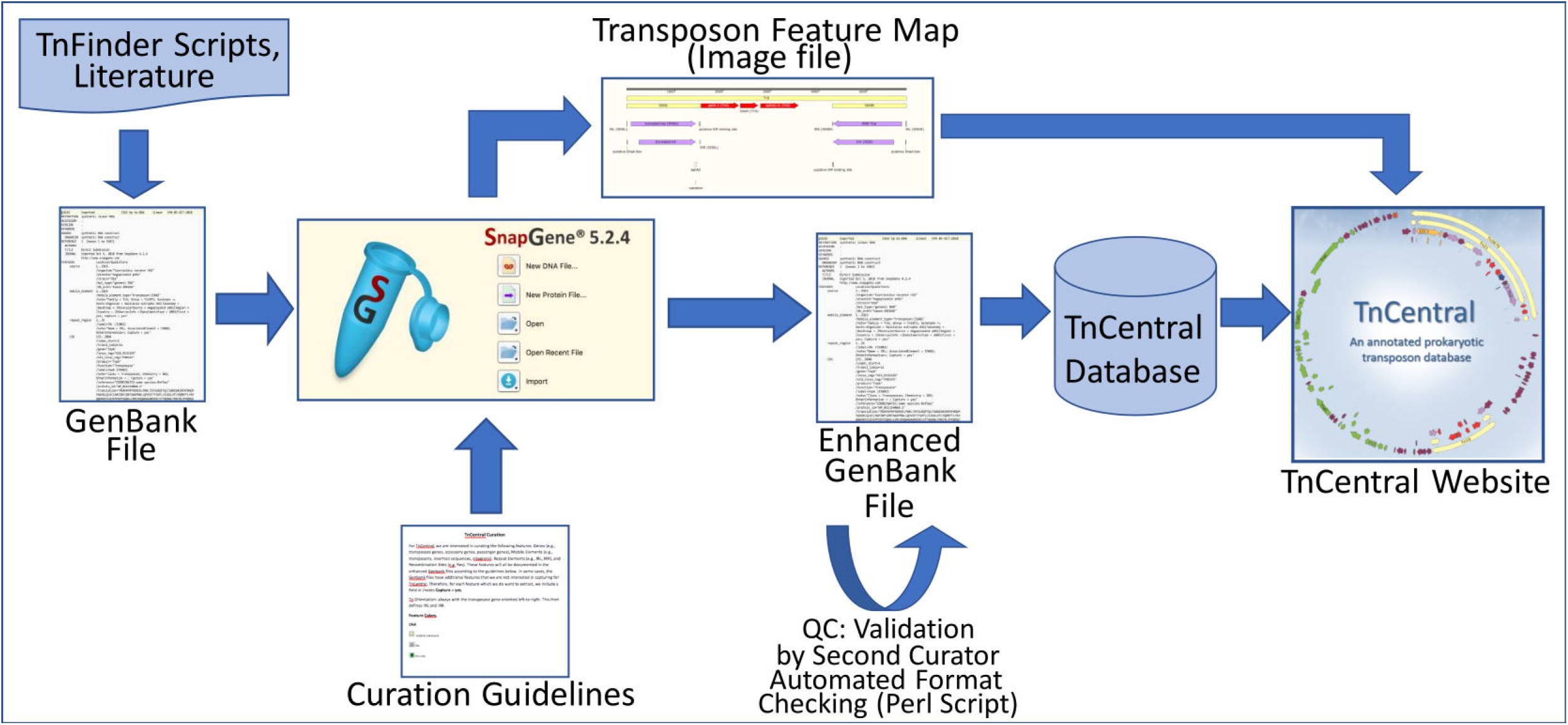
TnCentral Curation Workflow (see text for description).

Although we have adhered to the standard nomenclature for transposons extracted from the literature, for the many transposons newly identified during TnCentral database-building, we have temporarily used names indicating their source. In all cases, the Transposon Registry [53] accession number is provided as a synonym. There is some ambiguity in the literature concerning class 1 integrons and members of the Tn*402* transposon family. Class 1 integrons appear to be derivatives of this transposon family and include members with a range of Tn*402* transposition genes with varying degrees of completeness. We have therefore elected to include all Class 1 integrons as members of the Tn*402* family (Supplementary Table S1). ISfinder classification is used for the individual IS and in the case of compound transposons, the group to which they are belong is defined by the flanking IS.

Properties of protein coding genes are annotated with cross-references to database or ontology identifiers whenever possible. Antibiotic resistance gene properties, including gene name, sequence family, antibiotic resistance mechanism, and target drug classes are annotated according to the Antibiotic Resistance Ontology (ARO) as presented in Comprehensive Antibiotic Resistance Database (CARD) [10]. The Pfam [54] and InterPro resources [55] are used to define sequence family information.

### TnCentral Website implementation

TE features and sequence information are extracted from the enhanced GenBank files. TE feature information is used for the search and the entry pages, and the TE DNA and protein sequence information are used for the Sequence Search and display. The extracted data is loaded into the TnCentral database, implemented using MySQL. The website is built on a Linux server with Apache, and the web application is built on Perl CGI. Apache Lucene is used to index the data for flexible and fast search and retrieval. JavaScript is used for the interactive web-interface and display. BLAST is used for similarity search.

## Acknowledgements

We would like to thank John Dekker (NIAID, NIH Bethesda, Md, USA), Fred Dyda and Alison Hickman (NIDDK, NIH Bethesda, Md, USA), Patricia Siguier (CNRS Toulouse, France), Susu He (Nanjing University Medical School, Nanjing, China), Laurence van Melderen (Université Libre de Bruxelles, Belgium), and Gipsi Lima-Mendez and Bernard Hallet (Université de Louvain la Neuve, Belgium) for helpful discussions; Ben Glick (University of Chicago, SnapGene) for his help with the SnapGene software; the student curators at Georgetown University Medical Center for their contributions to the annotation process; and the Protein Information Resource (University of Delaware, Georgetown University Medical Center) for informatics support. This project was supported the U.S. Department of Defense (DoD) Global Emerging Infections Surveillance (GEIS) Branch (P0020_18_WR) and by institutional resources at the Center for Bioinformatics and Computational Biology, University of Delaware.

## Supplementary Figures

**Supplementary Figure 1.** Maps of Tn*5501* and Tn*5501.5* showing the loss of *parE* toxin gene in Tn*5501.5*. Feature color code: yellow--TE; purple--transposition genes; dark orange--toxin/anti-toxin genes; light orange--other open reading frames; grey--repeat elements; green--recombination sites. Maps were created with SnapGene.

**Supplementary Figure 2.** Maps of Tn*21* and its relatives. The feature color code is the same as in Supplementary Figure 1. Maps were created with SnapGene. Note that the different transposon derivatives are not to scale but their individual lengths are included.

**Supplementary Table S1.** The table displays the entire collection of TE at present in the database (May 2021) with columns indicating their **TnCentral accession numbers**, their **names, synonyms** from the literature and/or the Transposon Registry [53], TE **Type, Family Group**), **Host Organism** and **Molecular Source** (e.g., plasmid or chromosome). If no information is provided in the Molecular Source column, the source is chromosomal or unknown.

## Notes

https://tncentral.proteininformationresource.org/

### Competing Interest Statement

The authors have declared no competing interest.

